# Confounding by indication of the safety of de-escalation in community-acquired pneumonia: a simulation study embedded in a prospective cohort

**DOI:** 10.1101/652610

**Authors:** Inger van Heijl, Valentijn A. Schweitzer, C.H. Edwin Boel, Jan Jelrik Oosterheert, Susanne M. Huijts, Wendelien Dorigo-Zetsma, Paul D. van der Linden, Marc J.M. Bonten, Cornelis H. van Werkhoven

**Affiliations:** Department of Clinical Pharmacy, Tergooi hospital, Van Riebeeckweg 212, 1213 XZ Hilversum, The Netherlands; Julius Center for Health Sciences and Primary care, University Medical Centre Utrecht, Heidelberglaan 100, 3584 CH Utrecht, The Netherlands; Department of Medical Microbiology, University Medical Centre Utrecht, Heidelberglaan 100, 3584 CH Utrecht, The Netherlands; Department of Internal Medicine & Infectious Diseases, University Medical Centre Utrecht, Heidelberglaan 100, 3584 CH Utrecht, The Netherlands; Department of Pulmonary Medicine, University Medical Centre Utrecht, Heidelberglaan 100, 3584 CH Utrecht, The Netherlands; Department of Medical Microbiology, Tergooi hospital, Van Riebeeckweg 212, 1213 XZ Hilversum, The Netherlands

## Abstract

**Background:** Observational studies have demonstrated that de-escalation of antimicrobial therapy is independently associated with lower mortality. This most probably results from confounding by indication. Reaching clinical stability is associated with the decision to de-escalate and with survival. However, studies rarely adjust for this confounder. We quantified the potential confounding effect of clinical stability on the estimated impact of de-escalation on mortality in patients with community-acquired pneumonia.

**Methods:** Data were used from the Community-Acquired Pneumonia immunization Trial in Adults (CAPiTA). The primary outcome was 30-day mortality. We performed Cox proportional-hazards regression with de-escalation as time-dependent variable and adjusted for baseline characteristics using propensity scores. The potential impact of unmeasured confounding was quantified through simulating a variable representing clinical stability on day three, using data on prevalence and associations with mortality from the literature.

**Results:** Of 1,536 included patients, 257 (16.7%) were de-escalated, 123 (8.0%) were escalated and in 1156 (75.3%) the antibiotic spectrum remained unchanged. The adjusted hazard ratio of de-escalation for 30-day mortality (compared to patients with unchanged coverage), without adjustment for clinical stability, was 0.36 (95%CI: 0.18-0.73). If 90% to 100% of de-escalated patients were clinically stable on day three, the fully adjusted hazard ratio would be 0.53 (95%CI: 0.26-1.08) to 0.90 (95%CI: 0.42-1.91), respectively. The simulated confounder was substantially stronger than any of the baseline confounders in our dataset.

**Conclusions:** With plausible, literature-based assumptions, clinical stability is a very strong confounder for the effects of de-escalation. Quantification of effects of de-escalation on patient outcomes without proper adjustment for clinical stability results in strong negative bias. As a result, the safety of de-escalation remains to be determined.

## Introduction

The aim of antimicrobial stewardship is improving antibiotic use, without compromising clinical outcomes on the individual level (1). De-escalation of empirical antimicrobial therapy is highly recommended in antimicrobial stewardship programs. In a recent systematic review de-escalation of empirical antimicrobial therapy was associated with a 56% (95% CI 34%-70%) relative risk reduction in mortality (2). Although it seems a safe strategy, most studies evaluating de-escalation and reporting mortality were observational with a high risk of bias and with high clinical heterogeneity. As it seems highly unlikely that unnecessary broad-spectrum antibiotics lead to increased mortality in individual patients, the association between de-escalation and improved survival in observational studies is most likely biased by unmeasured confounding by indication. Confounding by indication is present if the indication for the intervention (here: de-escalation of empirical antimicrobial therapy) is also a prognostic factor for the outcome (mortality). De-escalation is usually only performed when clinical stability is reached in the first days after starting antimicrobial therapy and this also is a strong prognostic factor for patient outcome. However, hardly any of the observational studies adjusts for clinical stability during admission. In the aforementioned systematic review (2) only one of nineteen observational studies corrected for this confounder (3). Not taking this into account causes a negative bias (towards a protective effect). However, the magnitude of this bias has never been established. The aim of the current study was to quantify the potential effect of unmeasured confounding by indication (due to clinical stability) in the association between de-escalation and patient outcome in patients with community-acquired pneumonia.

## Methods

### Data collection

Data were used from the Community-Acquired Pneumonia immunization Trial in Adults (CAPiTA) (4). This study was a parallel-group, randomized, placebo-controlled, double blind trial to assess the efficacy of a 13-valent pneumococcal conjugate vaccine. The study included 84,496 immunocompetent community-dwelling adults, 65 years of age and above. Surveillance for suspected pneumonia was performed in 58 hospitals in the Netherlands, in the period September 2008 - August 2013. The study was approved by the Central Committee on Research Involving Human Subjects and by the Ministry of Health, Welfare and Sport in the Netherlands and all the participants provided written informed consent. For the current analysis, patients with a working diagnosis of CAP admitted to a non-intensive care unit (ICU) and receiving antibiotics on the day of admission were included. Patients were excluded from the current analysis if they participated in a simultaneously running interventional trial evaluating different antibiotic regimens for CAP (5), since this trial interfered with the choice of empirical antibiotic treatment, or if they died within 24 hours of admission because these are not eligible for de-escalation.

### Definitions

To define de-escalation, antibiotics were ranked based on their spectrum of activity against CAP pathogens, from rank 1 (‘narrow-spectrum’) to rank 3 (‘extended / restricted spectrum’) antibiotics (Table 1). In patients with combination therapy, the highest rank of any individual antibiotic was counted, except for combination therapy of β-lactam therapy and a macrolide, which was considered as rank 3, as for respiratory pathogens this combination results in a much broader spectrum than any of the individual antibiotics. Therapy adjustment was defined as the first switch from empirical therapy to another antimicrobial class during hospitalization, independent of the reason for switching. De-escalation and escalation were defined as a change to a lower rank or a higher rank, respectively. Continued regimens or adjustments to an equivalent rank were defined as continuation.

**Table 1:** Antibiotic ranking.

| **Rank 1**<br>**(Narrow spectrum)** | **Rank 2**<br>**(Broad spectrum)** | **Rank 3**<br>**(Extended / restricted spectrum)** |
| --- | --- | --- |
| Penicillin | 1<sup>st</sup> generation cephalosporins | 3<sup>d</sup> generation cephalosporins |
| Amoxicillin | 2<sup>nd</sup> generation cephalosporins | 4<sup>th</sup> generation cephalosporins |
| Tetracyclines | Co-amoxi-clav | Fluoroquinolones |
|  | Co-trimoxazole | Aminoglycosides |
|  | Clindamycine | Piperacillin/tazobactam |
|  | Macrolides | Carbapenems |
|  |  | Vancomycin |

### Statistical analysis

Descriptive statistics were used to describe clinical practice of de-escalation. Differences in patient characteristics between patients with a de-escalation versus no de-escalation were compared using Student’s *t* test or *χ*^2^ tests. Frequencies of de-escalation, escalation and continuation were described visually and numerically. We tested the proportional hazard assumptions for a follow-up period of 90 days, which revealed that the hazards were proportional up to 30 days and not thereafter (see Supplementary Figure 1). Therefore we used 30-day mortality as the outcome. To determine the effect of de-escalation on clinical outcome we excluded patients starting in rank 1, since they are not able to de-escalate. We performed Cox proportional hazards regression with de-escalation as time-dependent variable and adjusted for baseline characteristics using propensity score analyses. Propensity scores were calculated from a logistic regression model to estimate a patients propensity for de-escalation and included the variables: age, gender, smoking status, PSI-score, history of diabetes mellitus, history of chronic pulmonary disease, antibiotic use two weeks before admission, rank on day 1, season of admission, weekday vs. weekend day (the latter defined as Saturday or Sunday), and culture results. Propensity scores were then included as a continuous variable in the Cox proportional hazard regression model. Patients with escalation of therapy were censored at the time of escalation so that only the days before escalation contributed to the analysis. Other patients were censored at day 30.

### Effect of confounding by indication

To quantify the effect of unmeasured confounding by indication we simulated clinical stability during hospital admission as a new confounder. We defined clinical stability during admission as a binary variable evaluated at 72 hours, as therapy is often evaluated after three days. The strength of any given confounder is determined by the following three parameters: (1) the prevalence in the group with the determinant (de-escalation), (2) the prevalence in group without the determinant (continuation) and (3) the association with patient outcome (mortality). For the simulation of clinical stability at 72 hours we reviewed the literature for reasonable assumptions for the three parameters.

We assumed that 80% of CAP patients admitted to a non-ICU ward will be clinically stable at day three, based on three randomized controlled trials evaluating intravenous to oral switches in patients (6–8). As the prevalence of clinical stability in the total study population is a weighted average of the prevalence of clinical stability in the de-escalation and the continuation group, the prevalence in one group can be calculated from the prevalence in the other group. We assumed a high prevalence for clinical stability in the de-escalation group, so we varied the prevalence from 80% to 100%, with corresponding calculated prevalence’s in the continued group between 80% and 75% to arrive at the overall prevalence of 80%. The assumed crude odds ratio (OR) between clinical stability at 72 hours and 30-day mortality was 0.14, based on unpublished data of a randomized controlled trial evaluating the effect of adjunct prednisone therapy versus placebo on time to clinical stability for patients with CAP (Courtesy of dr. Blum) (9). In this trial, clinical stability was measured every 12 hours during hospital stay and was defined as time (days) until stable vital signs for ≥ 24 hours. To simulate the confounder of clinical stability at 72 hours in our dataset, we randomly assigned the presence and the absence of clinical stability such that the aforementioned assumptions about the three parameters were met. Subsequently, the HR of de-escalation on mortality adjusted for clinical stability was determined by including clinical stability as an extra covariate in the propensity score adjusted model. The consistency of the resulting adjusted HRs was tested by repeating the random assignment with a different random seed. In other words, the random assignment for the presence and absence of clinical stability was repeated three times, and resulted in the same adjusted HRs. In the end we plotted the crude and adjusted HR without clinical stability and the resulting HRs for different prevalence’s of clinical stability.

We also quantified the strength of each confounder as the change in HR of the model with or without each confounder. For the simulated confounder (clinical stability) we used the corresponding adjusted HR when added to the model with prevalence’s of resp. 90% and 100% in the de-escalation group. Data analysis was performed using SPSS for Windows, v.25.0 (SPSS, Chicago, IL, USA) and R v.3.4.3 https://www.R-projects.org/.

## Results

### Association between de-escalation and mortality

The study cohort consisted of 3,243 patients admitted with a clinical suspicion of pneumonia. After applying the in- and exclusion criteria 1,536 patients were included for analysis (Figure 1). Empirical treatment was rank-1 in 211 (13.7%), rank-2 in 624 (40.6%), and rank-3 in 701 (45.6%) patients. De-escalation occurred in 257 patients (16.7%) and escalation occurred in 123 (8.0%) patients. Most patients (1156, 75.3%) continued treatment without a change in rank of antimicrobial therapy during admission (Figure 2). Median time to de-escalation was 3.0 days (IQR 2.0 – 4.0 days). Compared to patients with continued (no de-escalation) regimens, patients with de-escalation less often were current smokers, more often had a pathogen identified and had a higher median PSI-score (Table 2). Patients in rank 2 de-escalated less often than patients in rank 3 (6.7% vs. 30.1%; p<0.001). Of the 257 patients with de-escalated therapy, therapy was later escalated in 14 patients (5.5%; 0.9% of all included patients) during admission.

**Figure 1:**
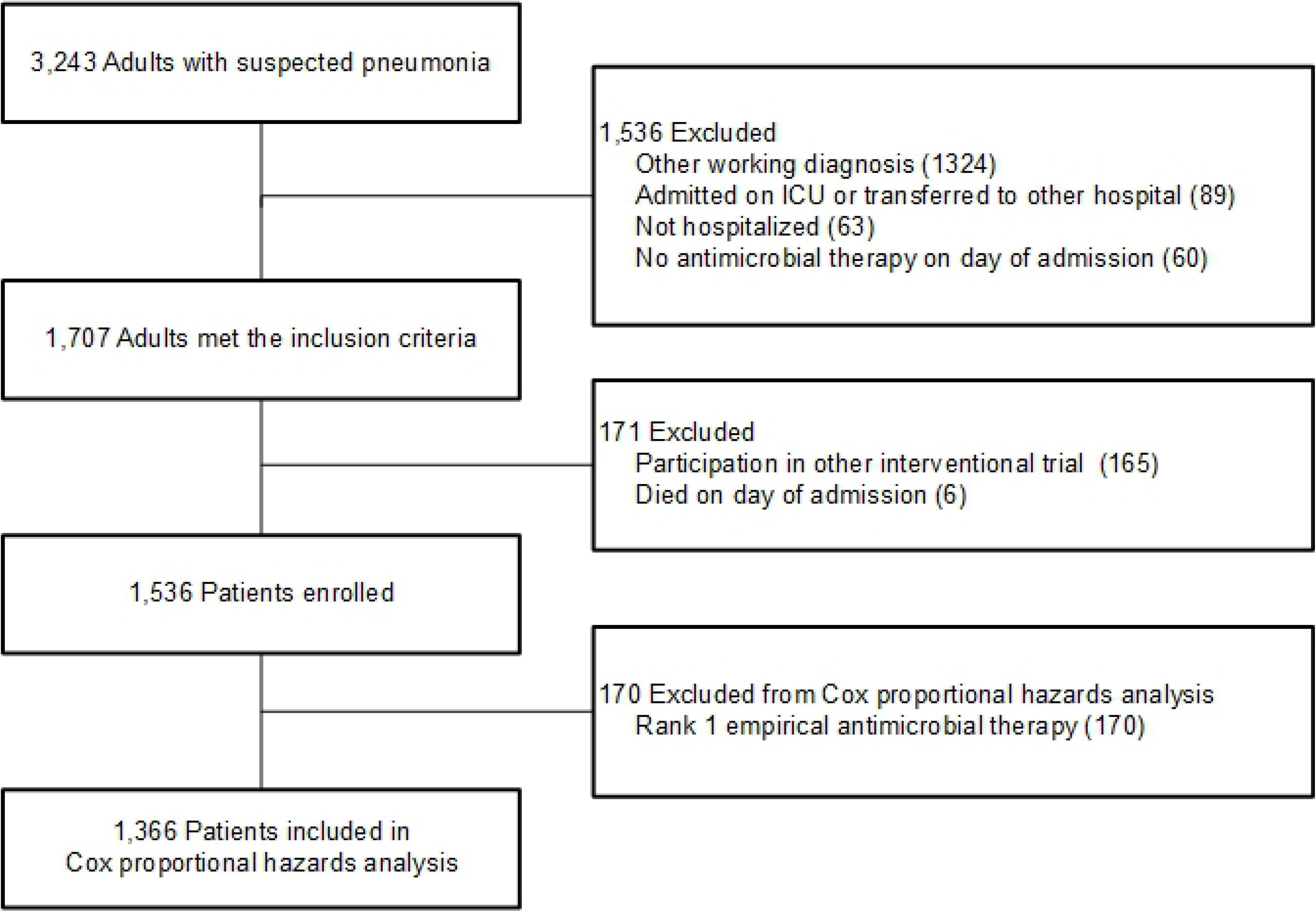
Patient selection.

**Figure 2:**
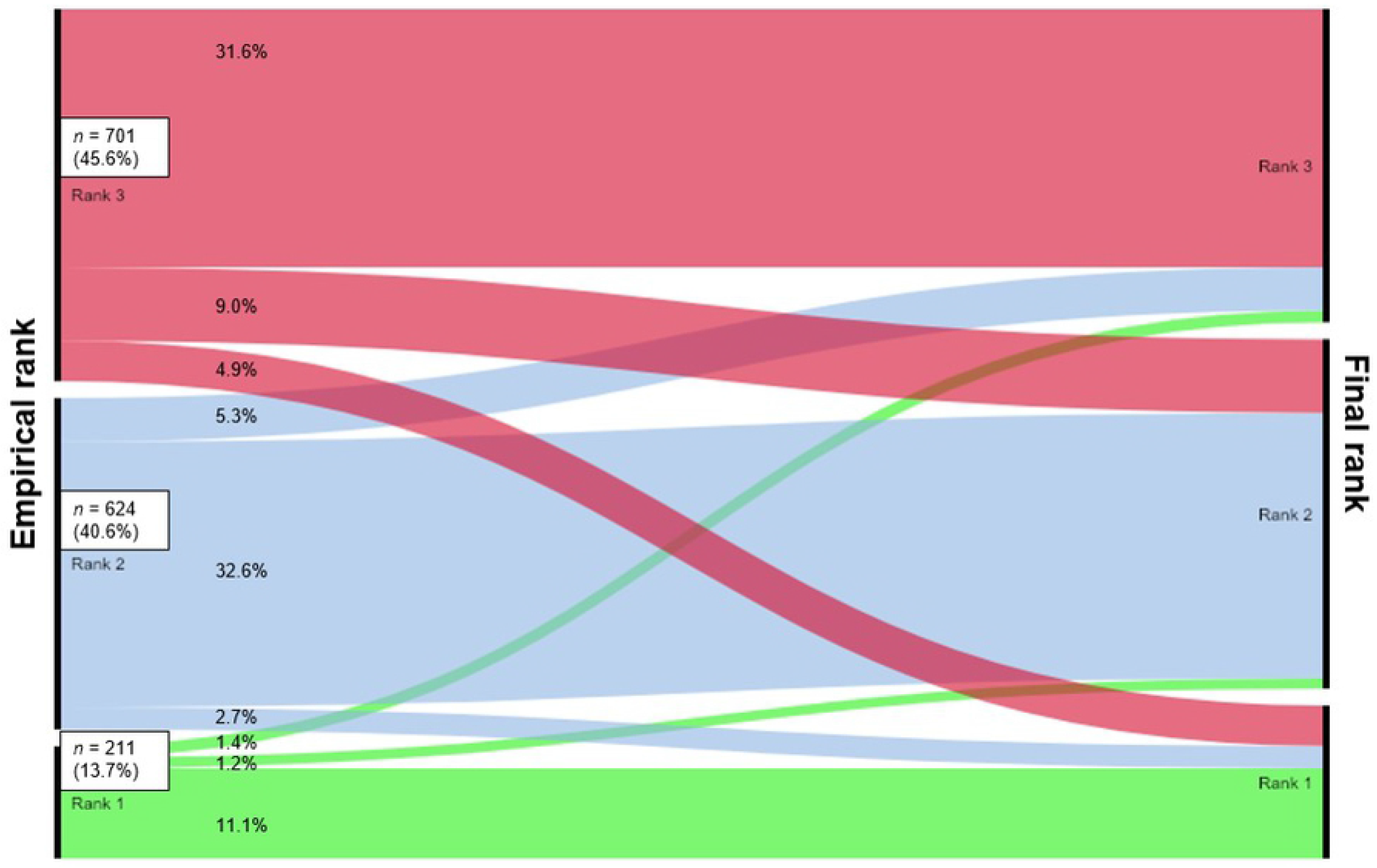
Alluvial-diagram of adjustment of empirical therapy. Rank-1 = narrow-spectrum-antibiotics, rank-2 = broad-spectrum-antibiotics, rank-3 = extended-spectrum-antibiotics. Continued regimen is a straight line, de-escalation is a falling line, escalation is a rising line.

**Table 2.** Baseline characteristics.

|  | Total cohort | De-escalation | No de-escalation (rank2-3)<sup>a</sup> | No de-escalation (rank 1)<sup>a</sup> |
| --- | --- | --- | --- | --- |
| Patients (N, %) | 1536 (100) | 257 (16.7) | 1068 (69.5) | 211 (13.7) |
| Age (y, median, range) | 77 (65 – 100) | 77 (66 – 95) | 77 (65 – 99) | 78 (66 – 100) |
| Male gender (n, %) | 1093 (71.2) | 189 (73.5) | 768 (71.9) | 136 (64.5) |
| Smoker (n, %) | 194 (12.6) | 21 (8.2) | 148 (13.9) | 25 (11.8) |
| Co-morbidities (n, %) |  |  |  |  |
| Chronic pulmonary disease | 849 (55.3) | 131 (51.0) | 608 (56.9) | 110 (52.1) |
| Chronic cardiovascular disease | 650 (42.3) | 123 (47.9) | 446 (41.8) | 81 (38.4) |
| Chronic renal disease | 11 (0.7) | 3 (1.2) | 5 (0.5) | 3 (1.4) |
| Chronic liver disease | 17 (1.1) | 5 (1.9) | 8 (0.7) | 5 (1.9) |
| Diabetes mellitus | 322 (21.0) | 53 (20.6) | 215 (20.1) | 54 (20.6) |
| PSI score (median, IQR) | 99 (82 – 117) | 103 (84-121) | 99 (82 – 118) | 92 (80 – 111) |
| Antibiotic use before admission (n, %) | 493 (32.1) | 84 (32.7) | 365 (34.2) | 44 (20.9) |
| Pathogen identified (n, %) | 469 (30.5) | 107 (41.6) | 303 (28.4) | 59 (28.0) |
| Day of admission (n, %) |  |  |  |  |
| Weekend | 613 (39.9) | 71 (27.6) | 263 (24.6) | 59 (28.0) |
| Empirical rank on day 1 (n, %) |  |  |  |  |
| Rank 1 | 211 (13.7) | NA | NA | 211 (100) |
| Rank 2 | 624 (40.6) | 42 (16.3) | 582 (54.5) | NA |
| Rank 3 | 701 (45.6) | 215 (83.7) | 486 (45.5) | NA |

Crude 30-day mortality was 3.5% (9/257) and 10.9% (107/986) in the de-escalation and continuation groups, respectively. The crude and adjusted hazard ratios for de-escalation, compared to continuation, were 0.40 (95% CI: 0.20 – 0.80) and 0.36 (95% CI: 0.18 – 0.73) for day-30 mortality. The AUC of the propensity score was 0.76 (95% CI: 0.73 – 0.79) and was considered acceptable.

### Effect of confounding by indication due to clinical stability

The results of the simulation analysis are depicted in Figure 3. Not using clinical stability for adjustment yields the afore-mentioned HR of 0.36. When using the assumed odds ratio between clinical stability at 72 hours and 30-day mortality of 0.14, the adjusted HR for de-escalation gradually increased to 0.90 with an increasing prevalence of clinical stability in patients with de-escalation up to 100%. Statistical significance of the adjusted HR was lost if the prevalence of clinical stability in the de-escalated patients was >= 89%. Determination of the strength of the simulated confounder, clinical stability, revealed that it was substantially stronger than any of the observed confounders in our dataset (Table 3).

**Figure 3:**
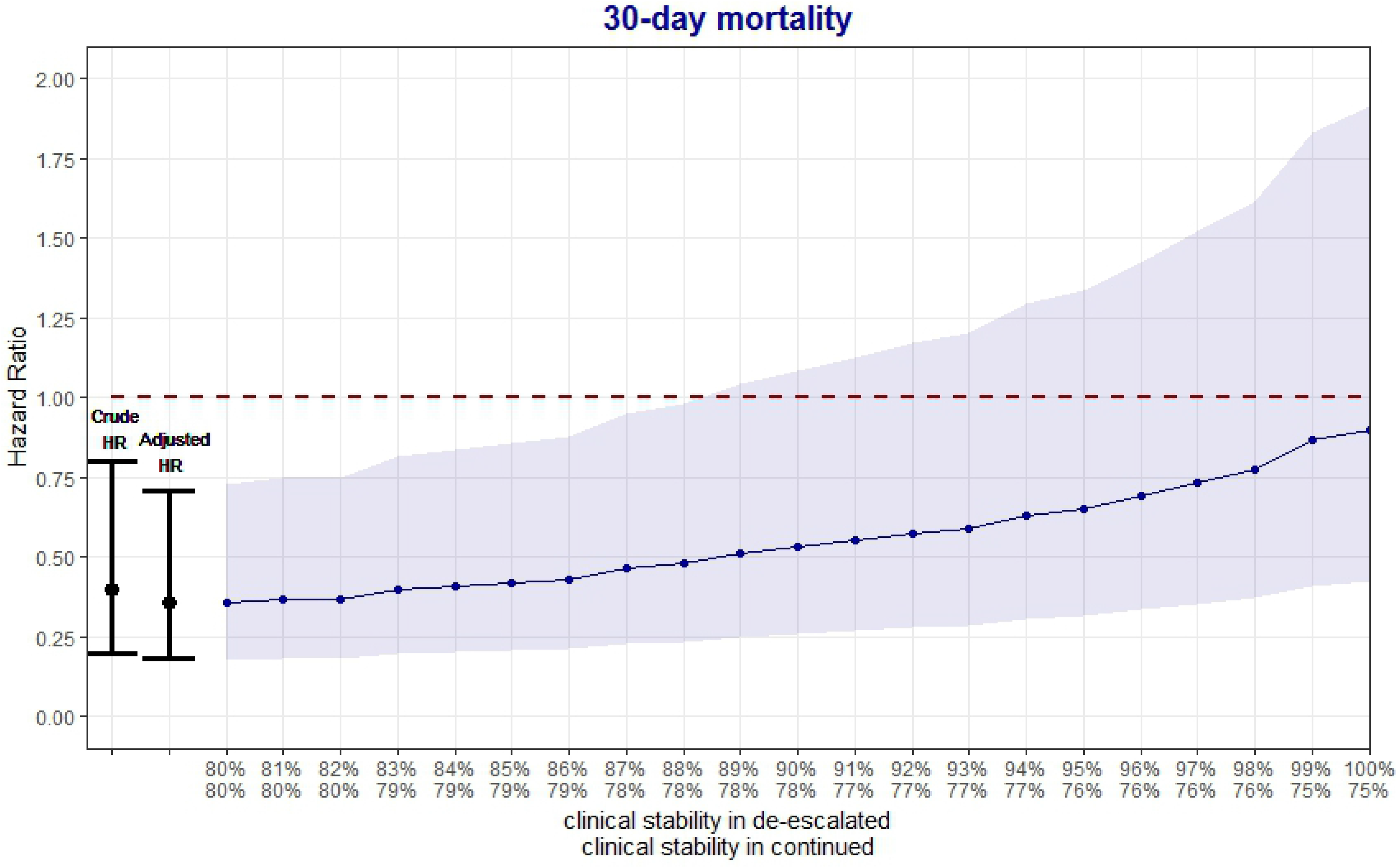
Effect of simulated confounder (clinical stability at 72 hours) on 30-day mortality. The line reflects the Hazard Ratios for 30-day mortality (based on Cox proportional hazard regression analysis adjusted with propensity scores) with 95% Confidence Interval (shaded area) for different prevalence’s of clinical stability in patients with and without de-escalation (horizontal axis). At the left side the weighted average of the two proportions is fixed at 80%, which reflects the adjusted Hazard Ratio without adjustment for clinical stability. The dashed line represents a HR of 1. The HR rises from 0.36 to 0.90 when the prevalence of clinical stability increases to 100% in the de-escalated group. Statistical significance is lost at a prevalence of clinical stability of 89% in the de-escalated group. For example a prevalence of 90% in de-escalated results in an adjusted HR of 0.53 (95% CI: 0.26 – 1.08) and a prevalence of 100% results in a HR of 0.90 (95% CI: 0.42 – 1.91).

**Table 3:** Strength of known and simulated confounders to the crude HR for 30-day mortality.

| **Confounder** | **% change of crude HR** |
| --- | --- |
| Diabetes mellitus | 0.1 |
| Antibiotic use before admission | 0.5 |
| Day of admission | 0.5 |
| Season of admission | 0.6 |
| Chronic pulmonary disease | 0.6 |
| Age | 0.9 |
| Culture results | 1.0 |
| Gender | 1.3 |
| Smoking | 1.5 |
| Rank on day 1 | 11.4 |
| Propensity score | 11.4 |
| PSI score | 14.8 |
| **Clinical stability (simulated)** |  |
| With prevalence in de-escalated group of 90% | 34.4 |
| With prevalence in de-escalated group of 100% | 122.7 |

## Discussion

In this observational study of patients hospitalized with CAP, after adjustment for observed baseline confounders de-escalation of antimicrobial therapy was associated with a 64% lower hazard of day-30 mortality. However, our simulations have demonstrated that clinical stability at 72 hours, which was not measured in our study, could fully explain this effect under reasonable, literature based assumptions. Based on these findings we conclude that the effects of de-escalation on patient outcome cannot be reliably quantified without adjustment for clinical stability, and that the true safety of de-escalation in antibiotic stewardship remains to be determined.

In a systematic review including different infectious diseases, de-escalation of empirical antimicrobial therapy was associated with a large reduction in mortality (2). Although our study only included CAP patients, we expect that the mechanism of bias applies to all infectious diseases for which empirical broad-spectrum antibiotic treatment is common practice. This bias, introduced by not including clinical stability during admission, applies to all previous studies evaluating de-escalation in patients with CAP hospitalized at a non-ICU ward (10–14). To the best of our knowledge, there are four observational studies on the association between de-escalation and mortality that adjusted for clinical stability or a similar time-varying confounder. In the first study by Joung et al. patients with intensive care unit-acquired pneumonia were included and clinical stability during admission was measured as two scores; APACHE-II (Acute Physiology and Chronic Health Evaluation II) and modified CPIS (clinical pulmonary infection score) both measured on day 5 after development of pneumonia. Both high APACHE II score (≥24) on day 5 and a high CPIS (≥10) on day 5 were associated with an increased 30-day pneumonia-related mortality. By including these confounders, next to other baseline covariates into the multivariable analysis the association between no de-escalation of antibiotics and 30-day mortality resulted in an aHR of 3.988 (95% CI 0.047 – 6.985) (15). The study objective was to determine independent risk factors for mortality, hence the focus was not on selecting appropriate confounders and one should be careful to interpret the results as a causal effect. Also, the wide confidence interval does not exclude a large increase in mortality in the de-escalated group. In the second study by Garnacho-Montero et al. patients admitted to the ICU with severe sepsis or septic shock were included and clinical stability during admission was measured as Sequential Organ Failure Assessment (SOFA) score on the day when culture results were available. A high SOFA score at culture result day was associated with a higher in-hospital mortality. When including this covariate next to other covariates the association between de-escalation and in-hospital mortality resulted in an aOR of 0.55 (95% CI 0.32 – 0.98, p = 0.022) (3). In the third study by Montravers et al. patients admitted with health care-associated intra-abdominal infection admitted to ICU were included and clinical stability during admission was measured by SOFA score. Here a decreased SOFA score at day three after initiation of empirical antimicrobial therapy was associated with a lower 28-day mortality. By including this covariate next to other covariates in the analysis this resulted in an aHR of 0.566 (95% CI 0.2503 – 1.278, p = 0.171) for association between de-escalation and 28-day mortality. However, this multivariate analysis also had the purpose to identify risk factors for 28-day mortality, not on selecting appropriate confounders (16). The fourth study by Lee et al. included patients with community-onset monomicrobial *Escherichia coli*, *Klebsiella* species and *Proteus mirabilis* bacteremia treated empirically with broad-spectrum beta-lactams and clinical stability during admission was measured by the Pitt bacteremia score. A high Pitt bacteremia score (≥4) at day three was associated with 4-week mortality. After propensity score matching there was no difference in crude mortality rates between de-escalation and no-switch regarding 2-week, 4-week and 8-week mortality (17). Comparison of the studies is difficult because different criteria for de-escalation and different definitions of disease severity during admission were used, and different populations were studied. A critique of the aforementioned studies is that all used de-escalation as a fixed variable. However, de-escalation is performed on a different day for each individual and should be analyzed as a time-dependent variable, otherwise it introduces immortal time bias (18).

It is recommended to include sensitivity analyses to estimate the potential impact of unmeasured confounding in every non-randomized study on causal associations (19). However, for observational studies evaluating de-escalation of antimicrobial therapy this has never been done before. To strengthen our sensitivity analysis we based our assumptions about the prevalence of clinical stability and association with mortality on existing high-quality data. We further assumed that physicians will only de-escalate when a patient is clinically stable or to initiate targeted treatment for an identified pathogen. In the latter case, we still expect that most patients in whom the physician decides to de-escalate will be clinically stable. We, therefore, expect that at least 90% and probably close to 100% of de-escalated patients will be clinically stable on day three.

Strengths of our study include the pragmatic approach of using prospectively collected data of a large patient population treated with empiric antibiotics and a working diagnosis of CAP. This included patients without an identified pathogen, which increases the generalizability of our study results. A limitation of our study is that we had to make assumptions for the prevalence of clinical stability in the de-escalated and continued group and for the association between clinical stability and day-30 mortality. These were derived from different study populations, all representing CAP patients hospitalized to a non-ICU ward. Our findings suggest that adjustment for clinical stability will result in a non-significant effect of de-escalation on mortality, which would be biologically plausible. Our findings also demonstrate that the individual baseline confounders, as measured in our study, only had minor effects on patient outcome, indicating that their correlation with clinical stability is rather weak. Another simplification in our analysis was that we modelled clinical stability as a binary variable on day 3, which does not well represent reality. For future studies we recommend to measure clinical stability repeatedly over time, as a time-varying confounder and on a continuous scale. Finally, we did not have information on quality of our sputum samples on which the pathogen was identified. Quality of sputum samples is also a prognostic factor for de-escalation of empirical antimicrobial therapy, however we could not correct for this in our model.

The results of our analysis also indicate that clinically relevant harm due to de-escalation cannot be excluded, as the upper boundary of the 95% confidence interval for the HR was close to 2 in the most extreme scenario. The scientific evidence for safety of de-escalation is de facto based on two RCTs. However, both RCTs are not powered for mortality. The first prospective, open-label, randomized clinical trial included patients with hospital-acquired pneumonia in an ICU without inclusion criteria regarding baseline clinical stability After randomization de-escalation was performed three to five days after initiation if empirical treatment when culture results were available. For the association between de-escalation and 14-day mortality the RR was 0.67 (95% CI 0.31 – 1.43), for 28-day mortality the RR was 0.75 (95%CI 0.46 – 1.23) and for in-hospital mortality the RR was 0.64 (95%CI 0.37 – 1.13), (calculated by the authors based on the data reported in (20). The other multicenter non-blinded randomized non-inferiority trial evaluated the safety of de-escalation with 90-day mortality as secondary outcome in patients with severe sepsis admitted to an ICU without inclusion criteria regarding baseline clinical stability. After randomization de-escalation was performed after culture results were available (IQR 2-4 days after initiation of empirical therapy). In the de-escalation group 18 of 59 patients (31%) died within 90-days, compared to 13 of 57 patients (23%) in the continuation group, yielding an adjusted HR of 1.7 (95% CI 0.79 – 3.49, p = 0.18). Although not statistically significant, this trend indicates potential harm rather than improved outcome due to de-escalation (21). As we have demonstrated, observational studies performed so far do not contribute to determining the safety of de-escalation because the amount of confounding by indication due to clinical stability is insurmountable. As appropriate adjustment of confounding by indication was not performed in the majority of the published observational studies on de-escalation, the ones that adjusted for clinical stability had other important limitations, and only two small RCTs have been performed, we conclude that the safety of this widely propagated antibiotic stewardship intervention remains unknown. We recommend that future observational studies addressing this research question include clinical stability in the analysis, preferably as a time-varying variable because clinical stability may change over time. Ultimately, although more expensive, de-escalation would be optimally studied in a pragmatic randomized controlled trial.

To conclude, the previously observed protective effect of de-escalation on mortality is likely due to confounding by unobserved factors such as clinical stability during admission.

## Acknowledgements

We gratefully acknowledge dr. Claudine Blum (Medizinische Universitätsklinik, Kantonsspital Aarau, Switzerland) and colleagues for sharing unpublished data and Lufang Zhang (University Medical Centre Utrecht, the Netherlands) for assistance with the data analysis.

